# Role of MicroRNA-22 (miR-22)/Specificity protein 1 (Sp1)/Cystathionine β-synthase (CBS) axis in regulating trophoblast invasion: an in vitro study

**DOI:** 10.1101/2023.03.08.531746

**Authors:** Pallavi Arora, Sankat Mochan, Sunil Kumar Gupta, Neerja Rani, Pallavi Kshetrapal, Sadanand Dwivedi, Neerja Bhatla, Renu Dhingra

**Affiliations:** All India Institute of Medical Sciences; All India Institute of Medical Sciences, New Delhi; cTranslational Health Science and Technology Institute

## Abstract

**Introduction:** Placenta expresses many ubiquitous as well as specific miRNAs which regulate trophoblast cell differentiation, proliferation, apoptosis, invasion, migration, and angiogenesis. Aberrant miRNAs expression (miR-22) has been linked to pregnancy complications, such as preeclampsia. Specificity protein 1 (Sp1) was identified a direct target of miR-22 and an inverse linear correlation between expression of miR-22 and Sp1 mRNA in gastric tumour has been reported. Also, Sp1 has a critical and indispensable role in the regulation of cystathionine β-synthase (H_2_S synthesizing enzyme). Exogenous H_2_S could promote cell proliferation and invasion by upregulating the expression of MMP-2 and MMP-9 in human bladder cancer EJ cells. Worldwide studies reported significant role of MMP-2 and MMP-9 in the regulation of trophoblast invasion.

**Aims and Objectives:** To determine the role of MicroRNA-22 (miR-22)/Specificity protein 1 (Sp1)/Cystathionine β-synthase (CBS) axis in regulating trophoblast invasion in the first-trimester trophoblast cells.

**Materials and methods:** The human first trimester trophoblast cell line (HTR-8/SVneo Cell line) was procured from American Type Culture Collection. Gain/Loss of function studies were conducted on HTR-8/SVneo cells and the expression of miR-22 (pre miR-22 and miR-22-3p), Sp1, CBS, MMP-2 and MMP-9 was observed. Invasion capacity of the cells was assessed by transwell invasion assay. Data were analyzed by STATA 14 and Graph Pad Prism 8.

**Results:** Significantly down regulated regulated mRNA and protein expression of Sp1, CBS and and MMPs 2, 9 was observed when the cells were transfected with miR-22-3p mimic in comparison to miR-22-3p inhibitor transfected cells. Expression of CBS and MMPs 2, 9 was found to be significantly up regulated when the cells were transfected with Sp1 plasmid as compared to Sp1 inhibitor (2,4,5 trifluoroaniline) treated cells at both transcription and translation levels. MMP-2 and MMP-9 levels were found to be significantly elevated when the cells were subjected to NaHS [sodium hydrogen sulphide (H_2_S donor)] treatment as compared to AOAA (aminooxyacetic acid) treated cells.

**Conclusion:** This is the first study which implies that miR-22 via targeting Sp1 must have influenced the mRNA and protein expression of CBS, MMP-2 and MMP-9 in order to regulate the trophoblast invasion.

## Introduction

MicroRNAs are small endogenous single-stranded RNAs that post transcriptionally regulate the expression of various target genes^1^. Worldwide studies suggest that miRNAs are important regulators of placental development^2,3,4–8^ and have been identified in human placental tissues^9^. Previous studies have shown that miRNAs regulate trophoblast cell proliferation, migration, invasion, apoptosis, and angiogenesis^4,7,8,10,11^ and their abnormal expression in placenta from women with compromised pregnancies like preeclampsia (PE) has been reported^3,4,12,13^. Specifically, miR-22 expression was found to be significantly up regulated in the placentae from pregnancies complicated by early onset preeclampsia as compared to preterm labor control placentae^14^. Specificity protein 1 (Sp1) has been recognized a direct target of miR-22^15^ and there is an inverse linear correlation between expression of miR-22 and Sp1 mRNA in gastric tumors has been observed. Also, miR-22 decreases CD147 (cluster of differentiation 147) expression through targeting Sp1 and the tumor suppressor like activity of miR-22 is due, in part, to down regulation of Sp1^16^. In Drosophila SL2 cells, co-transfection studies have been conducted which indicated that both human cystathionine β-synthase (CBS) gene promoters (−1a and -1b) are transactivated by Sp1 and Sp3 but only the -1b promoter is subject to a site-specific synergistic regulatory interaction between Sp1 and Sp3^17^. Hydrogen sulphide (H_2_S) is the third gas signaling molecule observed in the human body, which is generated by essential amino acid cysteine in vivo through one carbon unit metabolism and transfer-sulfur pathway with enzyme catalysis and the included enzymes were mainly cystathionine β-synthase (CBS), cystathionine γ-lyase, and 3-mercaptopyruvate sulfurtransferase^18-21^. Exogenous H_2_S [NaHS (sodium hydrogen sulphide)] was reported to promote cell proliferation and invasion by upregulating the expression level of matrix metalloproteinases (MMPs) 2, 9 in human bladder cancer EJ cells^22^. The invasion cascade exhibited by placental trophoblasts and cancerous cells bears many similarities, and it is attributed to extracellular matrix degradation mediated by MMPs^23^. Among MMPs, MMP-2 and MMP-9 appear to play a crucial role in regulating trophoblast invasion^24^ and their expression was found to be down regulated in preeclamptic and IUGR placentae^25^.

The aim of the present study was to determine the expression of miR-22, Sp1, CBS and MMPs 2, 9 in the first-trimester trophoblast cells and to discern whether miR-22 via targeting Sp1 regulates the expression of CBS and MMPs 2, 9 in HTR-8/SVneo cells at both transcription and translation levels.

## Methodology

The human first trimester trophoblast cell line (HTR-8/SVneo Cell line) was procured from American Type Culture Collection and maintained in Dulbecco’s Modified Eagle Medium and Ham’s F12 medium in 1:1 ratio and supplemented with 10% fetal bovine serum, 100U/ml penicillin, 100µg/ml streptomycin and amphotericin B 250 µg/ml at 37°C with 5% CO_2_ in tri-gas CO_2_ incubator (Thermo). Cells were passaged with 0.25% trypsin and 0.01% EDTA.

### Cell transfection and treatments

HTR/SVneo cells were cultured in 6-well plates and were grown to 80 % confluency. Cells were transfected with miR-22-3p mimic (hsa-miR-22-3p mirVana miRNA mimic, Ambion), miR-22-3p inhibitor (MISSION Synthetic miRNA Inhibitor, Sigma) and Sp1 plasmid (Thermo) using Lipofectamine [RNA iMAX transfection reagent (Thermo)] and Opti-MEM (Gibco). Cells were treated with Sp1 inhibitor [2,4,5 Trifluoroaniline (Sigma Aldrich)], H_2_S donor [Sodium hydrogen sulphide (NaHS), Cayman Chemical Company] and CBS inhibitor [amimooxyacetic acid (AOAA), Sigma Aldrich). Doses were standardized and are mentioned in supplementary file. After miR-22-3p mimic and inhibitor transfection, levels of miR-22-3p were observed by qRT-PCR and mRNA and protein expression of Sp1, CBS and MMPs 2, 9 was determined. After Sp1 plasmid transfection and its inhibitor treatment, the expression of Sp1, CBS and MMPs 2, 9 was observed at both transcription and translation levels. When the cells were subjected to NaHS and AOAA treatments, mRNA and protein expression of CBS and MMPs 2, 9 was observed. After all the transfection and treatment experiments, invasive capacity of cells was assessed by transwell invasion assay.

### Real-Time Quantitative Reverse Transcription polymerase chain reaction (qRT-PCR)

RNA isolation from cells (after various transfections and treatments) was done using miRNeasy Mini Kit, Qiagen (gene expression of pre miR-22, miR-22-3p) and TRIzol™ Reagent, Invitrogen, Thermo Fisher Scientific (mRNA expression of Sp1, CBS, MMPs 2, 9). The quality of RNA was examined by denaturing gel and quantity was measured on Micro-Volume UV/Visible Spectrophotometer (Thermo Fisher Scientific-NanoDrop TM 2000). c-DNA synthesis was done using miScript II RT Kit, Qiagen (gene expression of pre miR-22, miR-22-3p) and revert aid H-minus reverse transcriptase kit, Thermo Fisher Scientific (mRNA expression of Sp1, CBS, MMPs 2, 9). Quality of cDNA was checked on 0.8% agarose gel, visualized by ethidium bromide (EtBr) stain under UV and quantity was measured on Micro-Volume UV/Visible Spectrophotometer. cDNA was subsequently used for qRT-PCR (CFX96 Touch™ Real-Time PCR Detection System, BioRad). miScript SYBR Green PCR Kit, Qiagen was used with miScript Precursor Assay, Qiagen and miScript Primer Assay, Qiagen to determine the gene expression of pre miR-22 and miR-22-3p in HTR-8/SVneo cells. U6 small nuclear RNA was used as internal control. qRT-PCR reactions were carried out in 20 µl volume, including cDNA (template), SYBR Green (Thermo Fisher Scientific), forward and reverse primers (Sigma) and nuclease free water to amplify cDNA for determining the mRNA expression of Sp1, CBS and MMPs 2, 9 in HTR-8/SVneo cells. Glyceraldehyde-3-phosphate dehydrogenase (GAPDH) mRNA was used as internal control. Primers were designed by NCBI and confirmed by In-Silico PCR.

### Immunofluorescence (IF)

Coverslips were coated with poly-L-lysine for 1 h at room temperature. Coverslips were rinsed, dried and were sterilized under UV light. Cells were grown on coverslips and were subjected to various transfections and treatments. Cells were briefly rinsed in phosphate-buffered saline (PBS) followed by incubating the cells in 100% methanol (chilled at - 20°C) at room temperature for 5 min. Cells were subsequently washed with ice-cold PBS. Antigen retrieval was done at 95°C for 10 min followed by washing in PBS for 5 minutes. Cells were incubated for 10 min with PBS containing 0.1 % Triton X-100 followed by three times washing in PBS for 5 min. Subsequently cells were incubated in 1% BSA in PBST (PBS+ 0.1% Tween 20) for 30 min to block unspecific binding of the antibodies. Cells were incubated in primary antibodies [Sp1 (Merck) at a dilution of 1:50, CBS (Abcam) at a dilution of 1:200, MMP-2 (Abcam) at a dilution of 1:100 and MMP-9 (Abcam) at a dilution of 1:100] diluted in 1% BSA in PBST for overnight at 4°C followed by three times washing in PBS. Subsequently, cells were incubated in the secondary antibodies [Sp1, CBS, MMP-2, MMP-9 (FITC conjugated, Abcam)] diluted in 1% BSA in PBST for 1 h at room temperature in the dark followed by three times washing in PBS. Vector True VIEW™ Autofluorescence Quenching Kit, Vector laboratories was used following the manufacturers’ instructions to diminish unwanted autofluorescence from non-lipofuscin sources. Mounting was done by fluoroshield mounting media with DAPI (Abcam). Stained slides were observed under Nikon Eclipse Ti-S elements microscope using NiS-AR software (version 5.1).

### Western Blot (WB)

Protein extraction from cells (after various transfections and treatments) was done with RIPA buffer (Thermo Fisher Scientific) and protease inhibitor cocktail. Separating and stacking gels were prepared. 3x non-reducing sample buffer was added to each of isolated protein sample from cells (after various transfections and treatments) followed by denaturation at 95°C for 5 minutes and the samples were loaded along with protein molecular marker (Thermo Fisher Scientific) to the wells. Subsequently, running the gel at 50 V (vertical electrophoresis apparatus, BioRad) in electrophoresis buffer for 3-3.5 hours until good band separation is achieved. Nitrocellulose membrane was used for the transfer of gel products on to the membrane in transfer buffer for 90 minutes. Membrane was washed with TBST20 followed by blocking in 5% BSA TBST20 for 90 minutes. Overnight incubation was done with primary antibodies [Sp1 (Merck) at a dilution of 1:400, CBS (Abcam) at a dilution of 1:1000, MMP-2 (Abcam) at a dilution of 1:1000 and MMP-9 (Abcam) at a dilution of 1:1000] at 4°C. Washing was done with TBST20 followed by incubation with secondary antibodies (Abcam) for 3 hours and then washed with TBST20. ECL kit (Thermo Fisher Scientific) was used for visualization of bands in Densitometer (Protein Simple).

### Transwell invasion assay (TIA)

100 µl diluted matrigel (Corning) was added in the insert (Greiner Bio-One, Germany) and incubated at 37°C overnight for gelling. 750 µl culture medium (DMEM, Ham’s F-12 medium with 10% FBS) was added in lower chamber and approximately 2.5 × 10^4^ cells (after various transfections and treatments) in serum free medium (DMEM and Ham’s F-12 medium) were placed on transwell insert (pore size 8.0 µm) followed by incubation at 37°C for 16 hours. Subsequently, medium was removed followed by washing with PBS and cells were fixed by formaldehyde (3.7% in PBS). Permeabilization was done by 100% methanol for 20 min followed by PBS washing. Staining was done by crystal violet for 15 minutes at room temperature. Subsequently, washing was done by PBS followed by scraping off non-invasive cells with cotton swabs and invasive cells were counted under Nikon Eclipse Ti-S microscope and DS-Fi2 camera. The number of invaded cells were counted on each transwell insert under 20x objective and sixteen fields were analyzed for each insert.

The percentage invasion and invasion index were calculated with the following formulae:

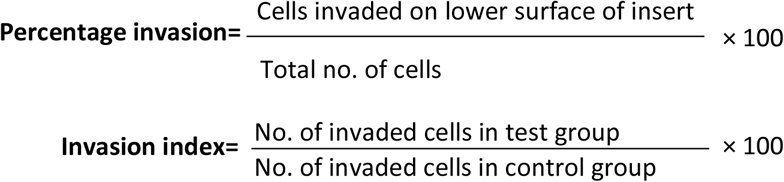

### Statistical Analysis

Data was analyzed by STATA 14 and Graph Pad Prism 8. Relative quantification cycles of gene of interest (ΔCq) were calculated by ΔCq = Cq (target) - Cq (reference). Relative mRNA expression with respect to internal control gene was calculated by 2^-ΔCq^. Paired t-test and wilcoxon matched-pairs signed rank tests were used to compare the average level of the variable between two groups and for more than two groups, one way ANOVA with Bonferroni’s post hoc test and Kruskal Wallis with Dunn test were applied. *p* value<0.05 was considered statistically significant.

## Results

The basal-level gene expression of miR-22-3p, pre miR-22, Sp1, CBS and MMPs 2, 9 was observed in HTR-8/SVneo cells (Figure 1).

**Figure 1.**
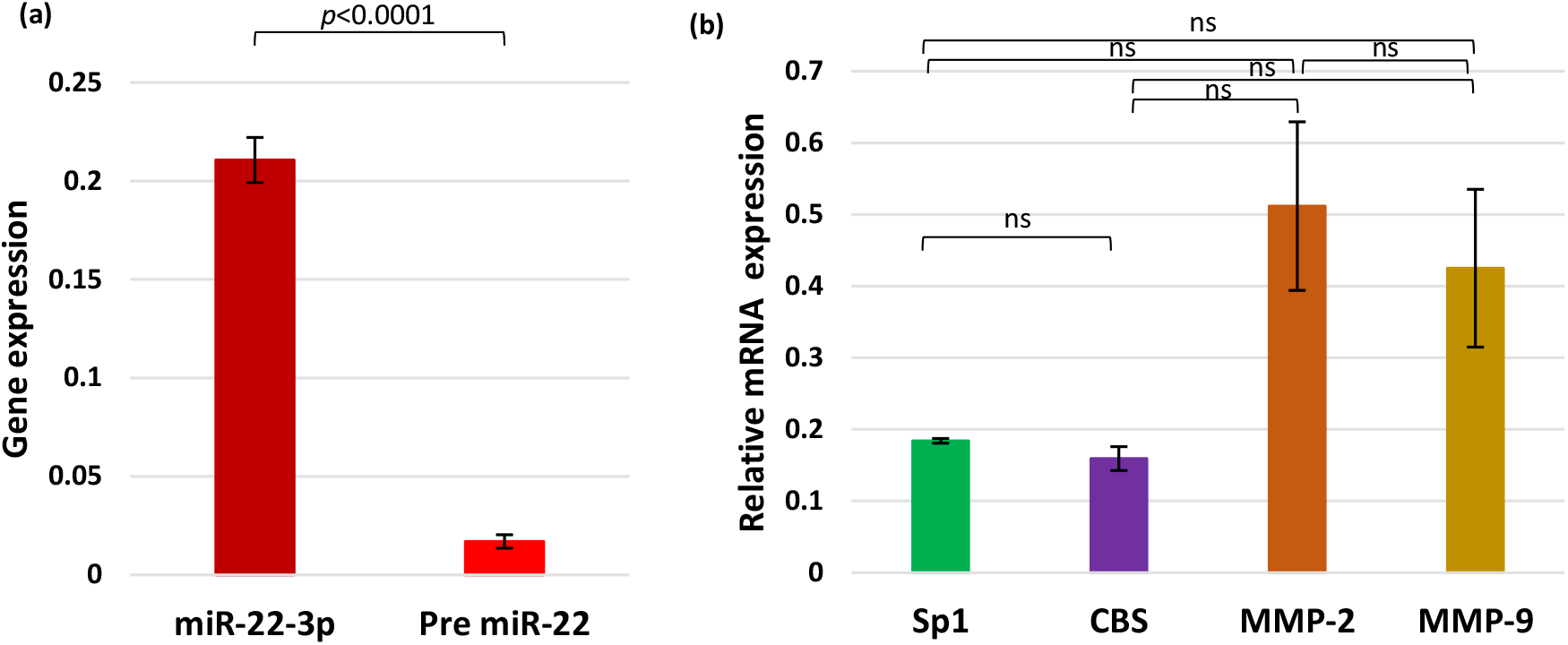
(a,b): Bar diagrams represent gene expression of pre miR-22, miR-22-3p, Sp1, CBS and MMPs 2, 9 in HTR-8/SVneo cells when no treatment was given. Experiments were performed three times in triplicate (n=9). Data presented as mean ± SEM. Paired t (a) and Kruskal Wallis with Dunn (b) tests were applied, ns: not significant.

**Figure 2:**
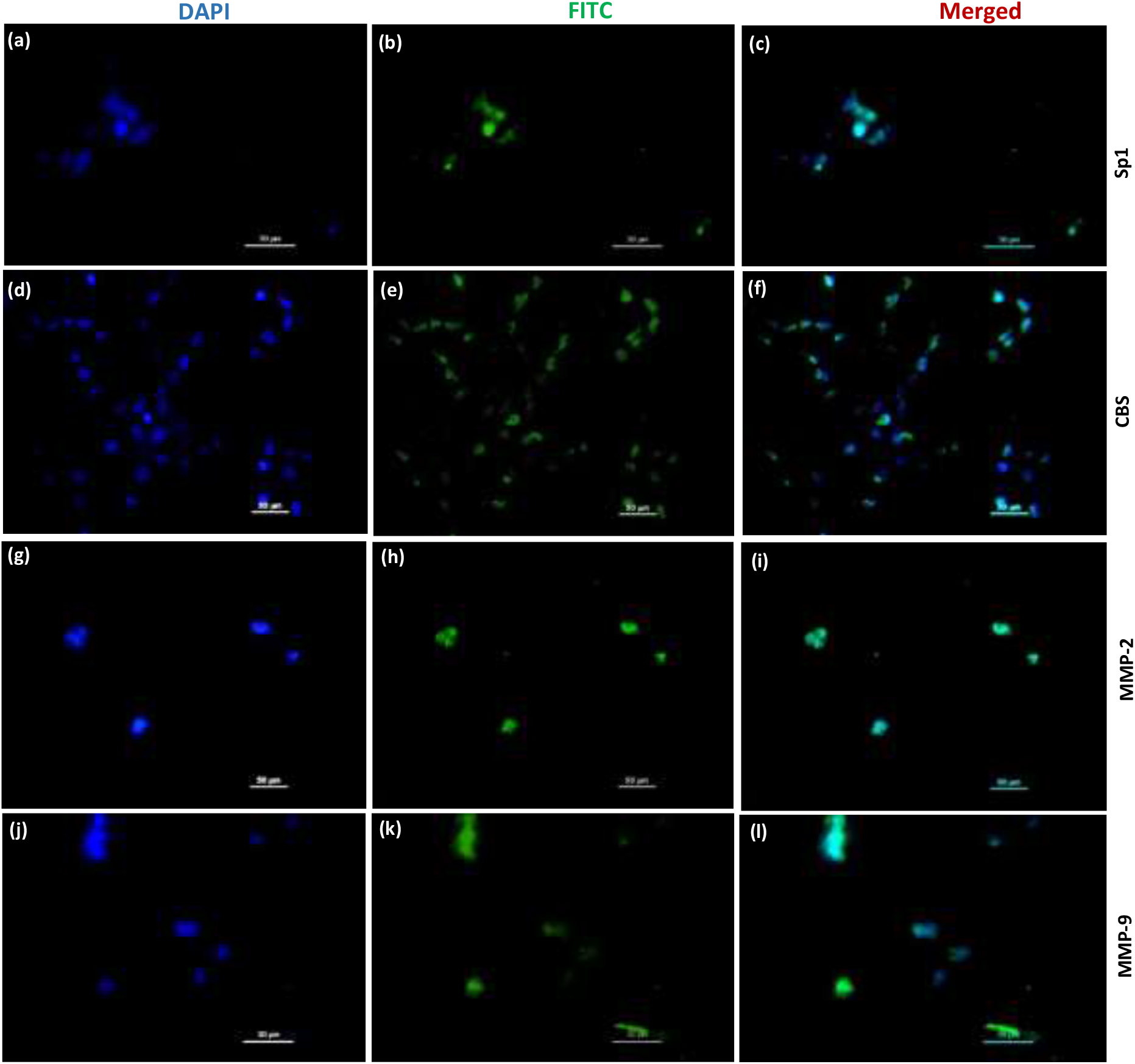
Representative photomicrographs of HTR-8/SVneo cells showing Sp1 (a,b,c), CBS (d,e,f), MMP-2 (g,h,i) and MMP-9 (j,k,l) protein expression when HTR-8/SVneo cells were given no treatment, a,d,g,j: DAPI: Blue, nuclear stain, b,e,h,k: FITC: Green, Primary antibodies Sp1, CBS, MMP-2, MMP-9 (secondary antibody conjugated with FITC), c,f,i,l: Merged images of DAPI and FITC. Scale bar: 50 µm

**Figure 3:**
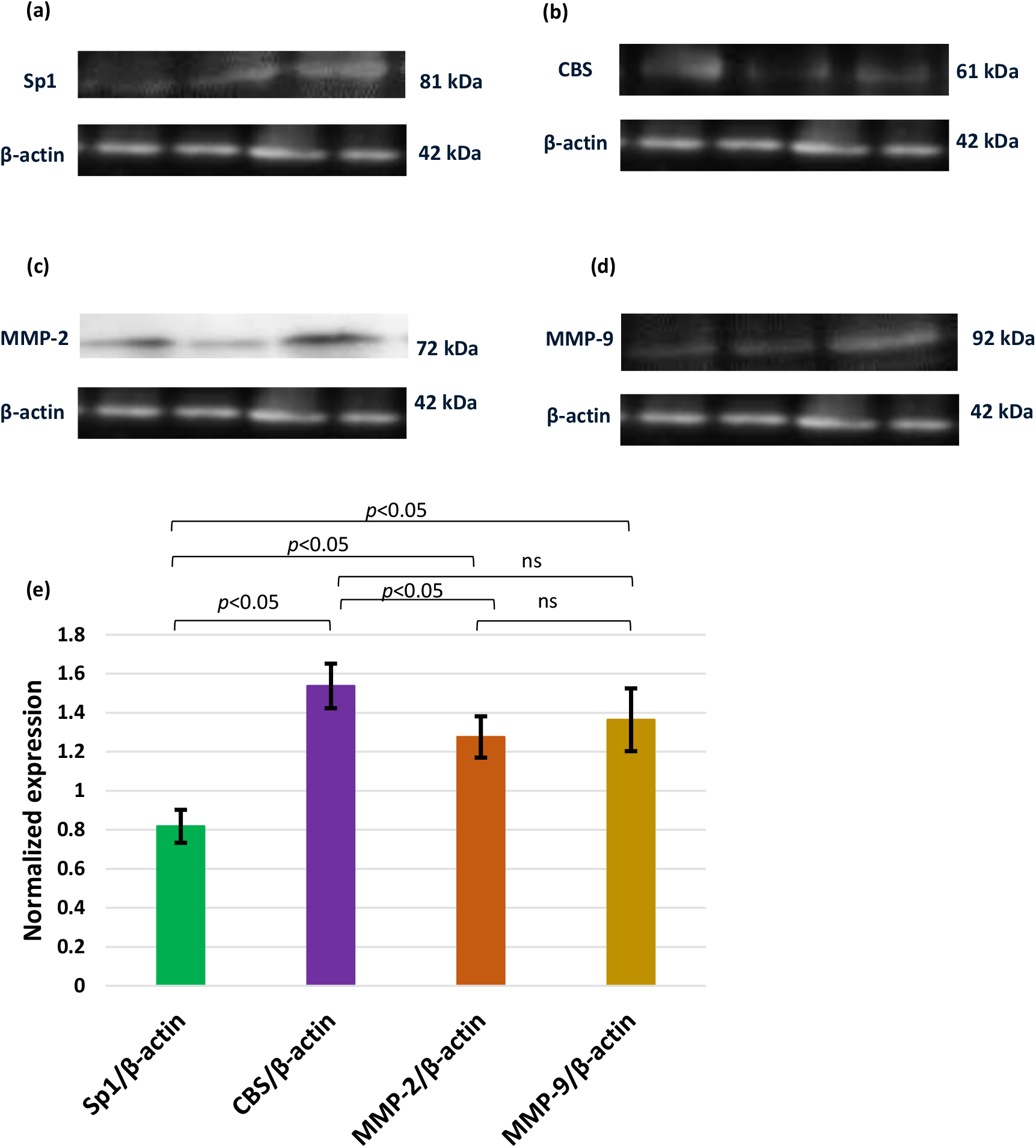
Representative images of immunoblot showing the protein expression of Sp1 (a), CBS (b), MMP-2 (c) and MMP-9 (d) when HTR-8/SVneo cells were given no treatment. Bar diagrams represent normalized expression of Sp1, CBS and MMPs 2, 9 proteins when HTR-8/SVneo cells were given no treatment (e). Experiments were performed three times in triplicate (n=9). Data presented as mean ± SEM. Kruskal Wallis with Dunn test was applied, ns: not significant.

Gene expression of miR-22-3p was found to be significantly up regulated as compared to pre miR-22 (Figure 1a). Therefore, cells were transfected with miR-22-3p mimic and miR-22-3p inhibitor to determine the gene expression of miR-22-3p and the mRNA and protein expression of Sp1, CBS and MMPs 2, 9.

Transfection of HTR-8/SVneo cells with miR-22-3p mimic leads to up regulation of the gene expression of miR-22-3p and when the cells were transfected with miR-22-3p inhibitor, gene expression of miR-22-3p got down regulated by qRT-PCR (Figure 4a).

**Figure 4:**
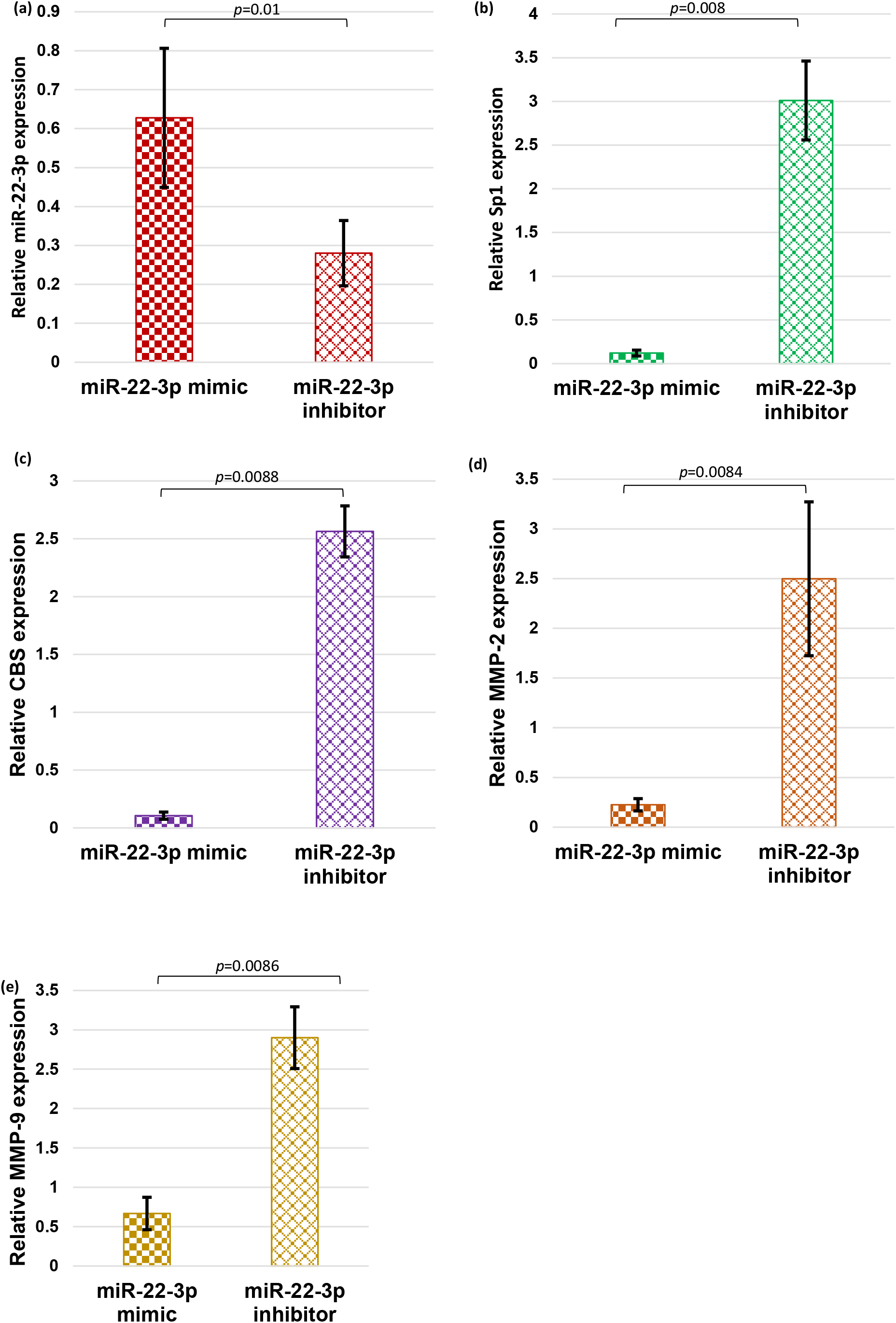
Bar diagrams represent gene expression of miR-22-3p, Sp1, CBS and MMPs 2, 9 post miR-22-3p mimic and inhibitor transfection. Experiments were performed three times in triplicate (n=9). Data presented as mean ± SEM. Wilcoxon matched-pairs signed rank test was applied.

### Transfection of HTR-8/SVneo cells with miR-22-3p mimic and miR-22-3p inhibitor

qRT-PCR data revealed significantly reduced mRNA expression of Sp1, CBS, MMP-2 and MMP-9 when HTR-8/SVneo cells were transfected with miR-22-3p mimic however, significantly elevated mRNA expression of Sp1, CBS, MMP-2 and MMP-9 was observed when the cells were transfected with miR-22-3p inhibitor [Figure 4 (b-e)]. IF staining demonstrated weak expression of Sp1, CBS, MMP-2 and MMP-9 proteins when the cells were transfected with miR-22-3p mimic but when the cells were transfected with miR-22-3p inhibitor, strong signals were observed for Sp1, CBS, MMP-2 and MMP-9 proteins (Figure 5). WB data showed significantly down regulated levels of Sp1, CBS, MMP-2 and MMP-9 proteins when the cells were transfected with miR-22-3p mimic whereas after miR-22-3p inhibitor transfection, protein levels of Sp1, CBS, MMP-2 and MMP-9 were found to be significantly up regulated (Figure 6). Transwell invasion assay analysis showed significant decrease in the number of invaded cells, percentage invasion and invasion index when the cells were transfected with miR-22-3p mimic whereas after miR-22-3p inhibitor transfection, the number of invaded cells, percentage invasion and invasion index were found to be significantly higher (Figure 7).

**Figure 5:**
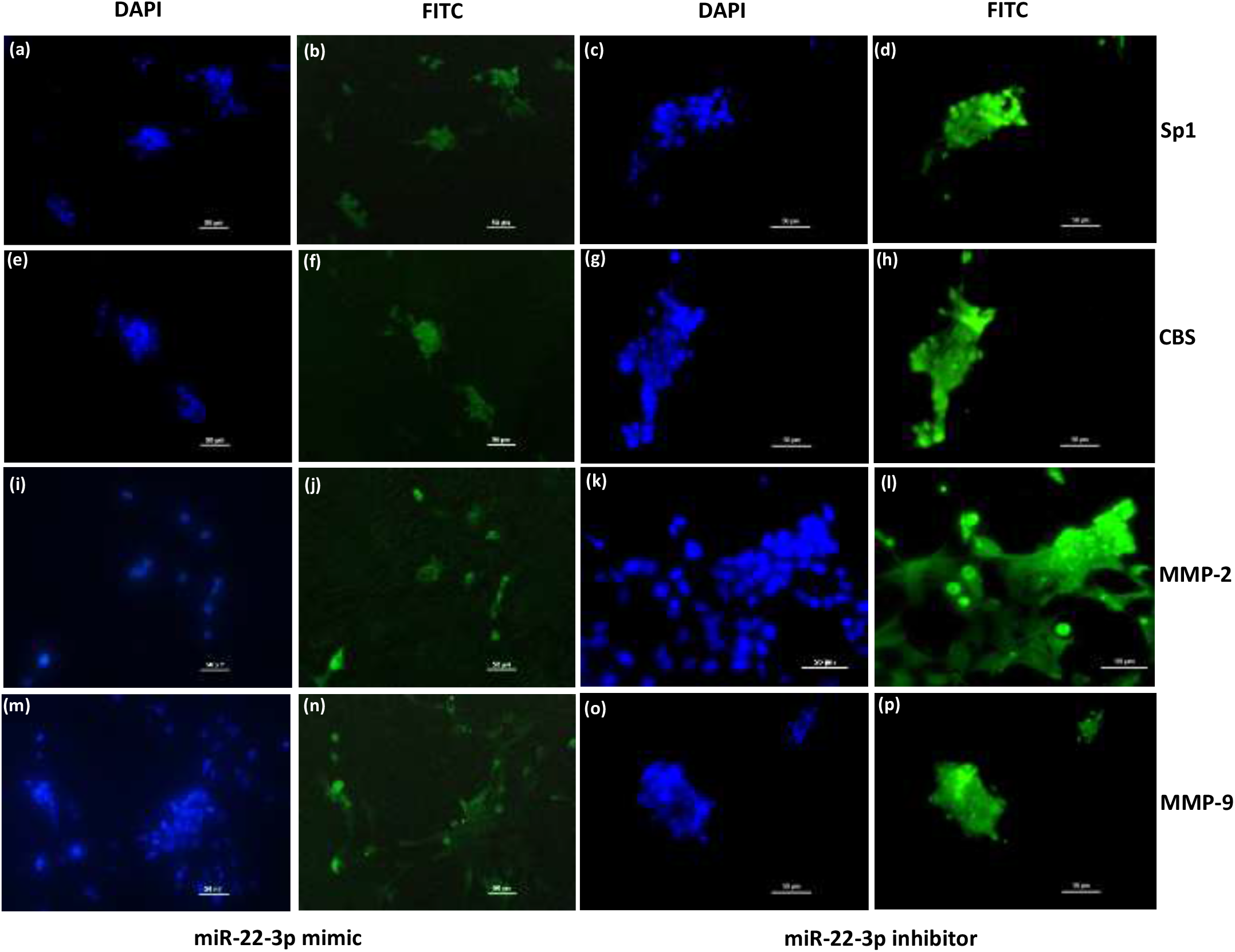
Representative Immunofluorescence images showing localization of Sp1 (b,d), CBS (f,h), MMP-2 (j,l) and MMP-9 (n,p) proteins stained with FITC when HTR-8/SVneo cells were transfected with miR-22-3p mimic and inhibitor. Nuclei were stained by DAPI (a,c,e,g,i,k,m,o); 20x magnification; scale bar: 50 µm.

**Figure 6:**
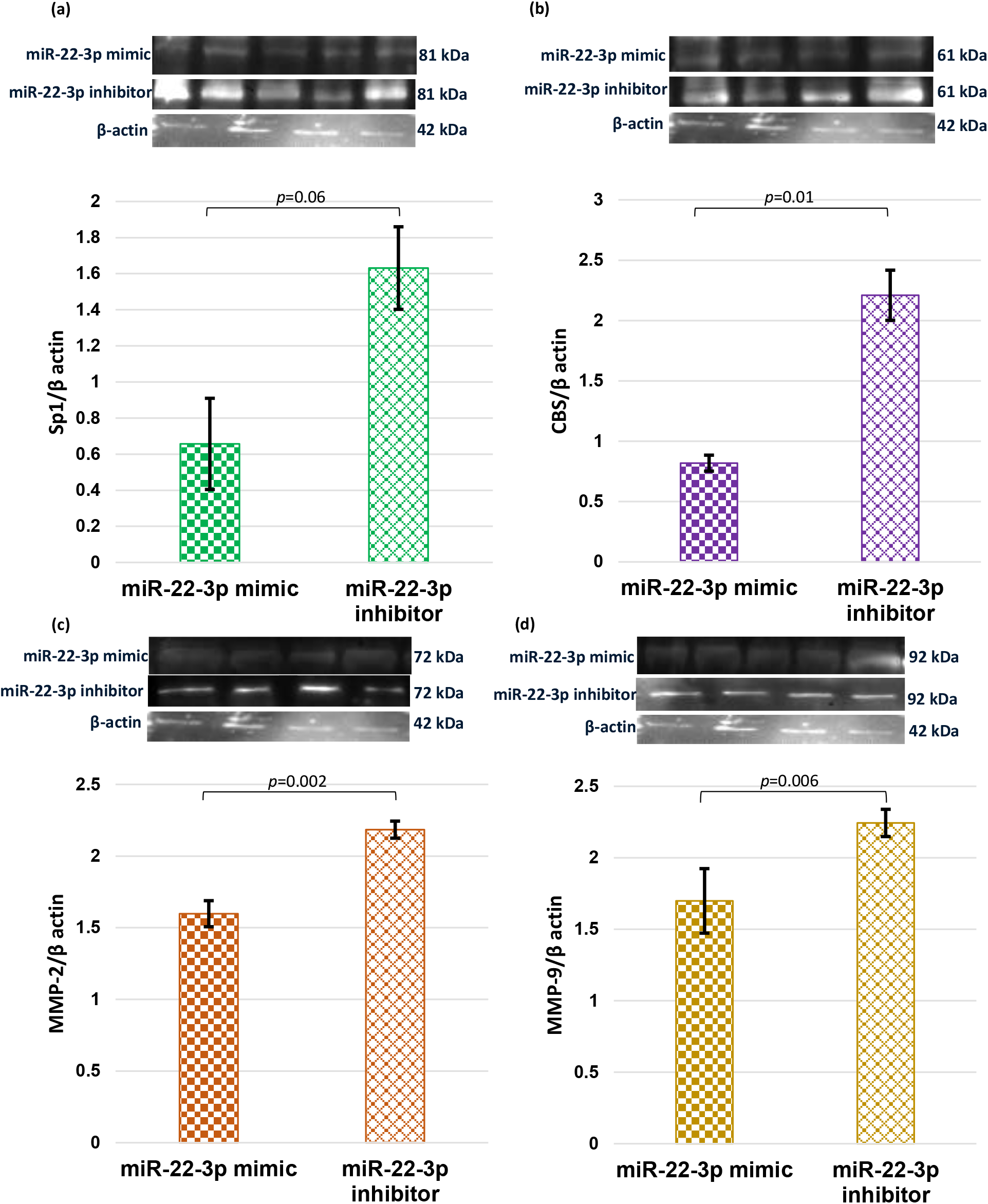
Representative images of immunoblot showing the protein expression of Sp1 (a), CBS (b), MMP-2 (c) and MMP-9 (d) when HTR-8/SVneo cells were transfected with miR-22-3p mimic and inhibitor. Bar diagrams represent the normalized values of Sp1 (a), CBS (b), MMP-2 (c) and MMP-9 (d) with respect to β-actin (loading control) post miR-22-3p mimic and inhibitor transfection. Experiments were performed three times in triplicate (n=9). Data presented as mean ± SEM. Statistical analysis was done using wilcoxon matched-pairs signed rank (Sp1) and paired t tests (CBS, MMPs 2, 9).

**Figure 7.**
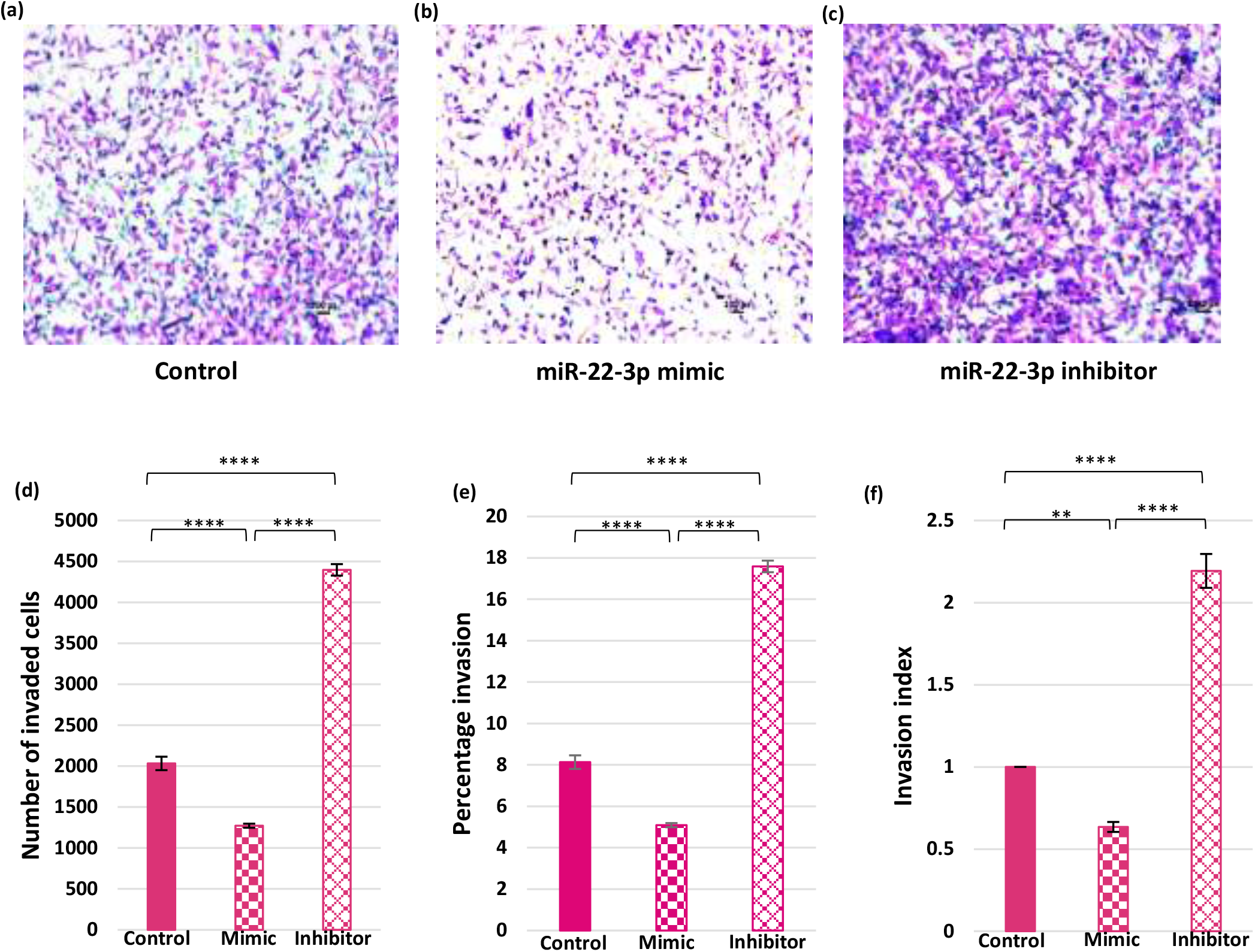
(a-c): Effect of miR 22-3p mimic and inhibitor transfection on the invasion capacity of HTR-8/SV neo cells. Representative matrigel transwell invasion assay images showing invaded cells on the underside of the membrane when no treatment was given (a) and after transfection with miR-22-3p mimic (b) and inhibitor (c), scale bar: 50 µm. 16 fields were used to count cells on each transwell insert. Bar diagrams represent number of invaded cells (d), percentage invasion (e) and invasion index (f) among various experimental groups. Experiments were performed three times in triplicate (n=9). Data presented as mean ± SEM, one way ANOVA with Bonferroni’s post hoc test was applied. ** *p*<0.01, *** *p*<0.001, \*\*\*\**p*<0.0001

### Transfection of HTR-8/SVneo cells with Sp1 mimic and treated with 2,4,5 Trifluoroaniline (Sp1 inhibitor)

qRT-PCR data revealed significantly elevated mRNA expression of Sp1, CBS, MMP-2 and MMP-9 when HTR-8/SVneo cells were transfected with Sp1 expression construct, however, significantly reduced gene expression of Sp1, CBS, MMP-2 and MMP-9 was observed when the cells were transfected with 2,4,5 Trifluoroaniline (Sp1 inhibitor) (Figure 8). IF staining demonstrated enhanced expression of Sp1, CBS, MMP-2 and MMP-9 proteins when the cells were transfected with Sp1 expression construct but when the cells were treated with 2,4,5 Trifluoroaniline, feeble expression was observed for Sp1, CBS, MMP-2 and MMP-9 proteins (Figure 9). WB data showed significantly up regulated protein levels of Sp1, CBS, MMP-2 and MMP-9 when the cells were transfected with Sp1 expression construct but when the cells were subjected to 2,4,5 Trifluoroaniline, protein levels of Sp1, CBS, MMP-2 and MMP-9 were found to be significantly down regulated (Figure 10). Transwell invasion assay analysis observed significant increase in the number of invaded cells, percentage invasion and invasion index when the cells were transfected with Sp1 mimic but when the cells were treated with 2,4,5 Trifluoroaniline, significant decrease in the number of invaded cells, percentage invasion and invasion index was reported (Figure 11).

**Figure 8:**
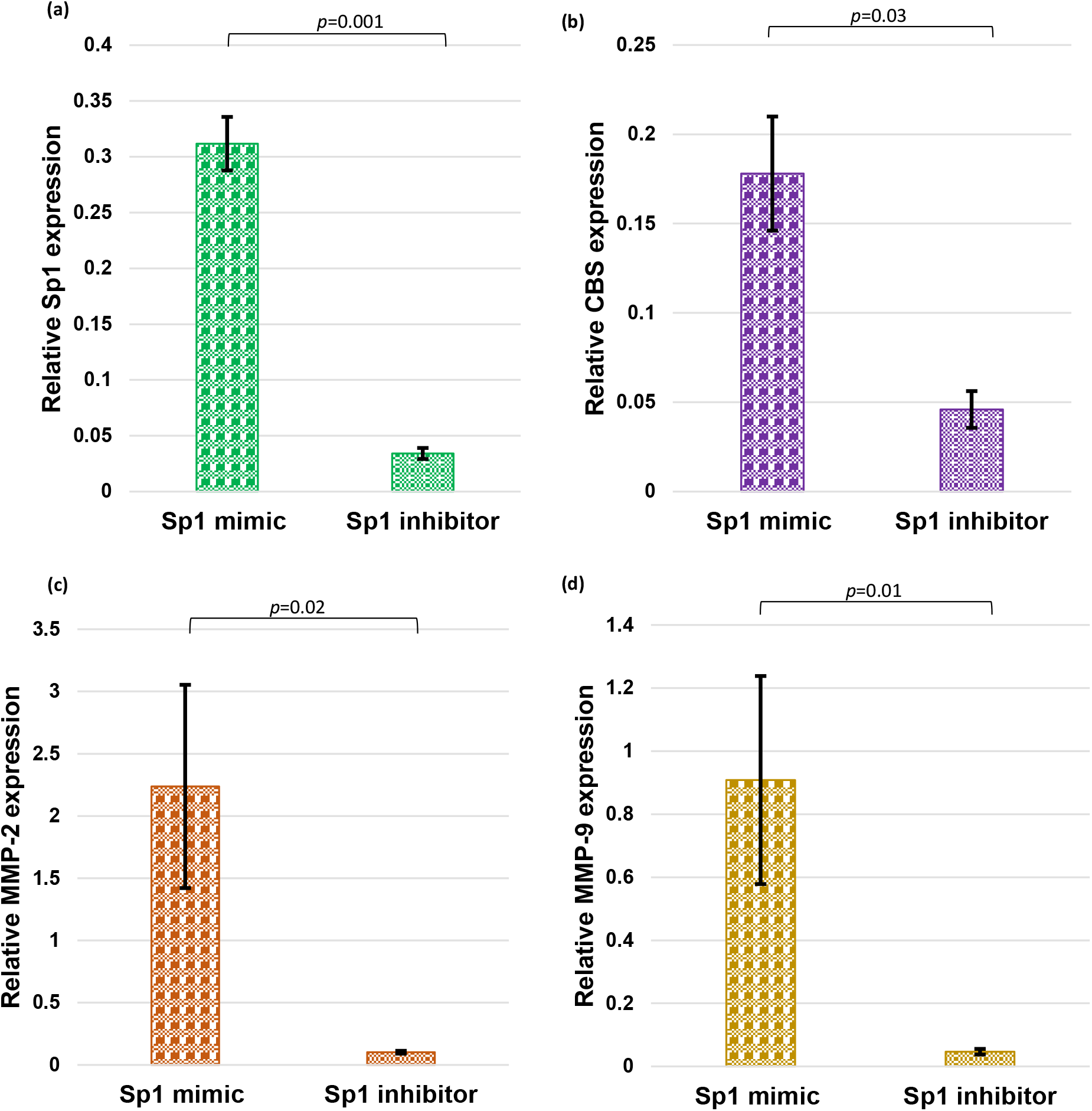
Bar diagrams represent gene expression of Sp1, CBS and MMPs 2, 9 when cells were transfected with Sp1 mimic and its inhibitor. Experiments were performed three times in triplicate (n=9). Data presented as mean ± SEM. Paired t (a) and wilcoxon matched-pairs signed rank (b-d) tests were applied.

**Figure 9:**
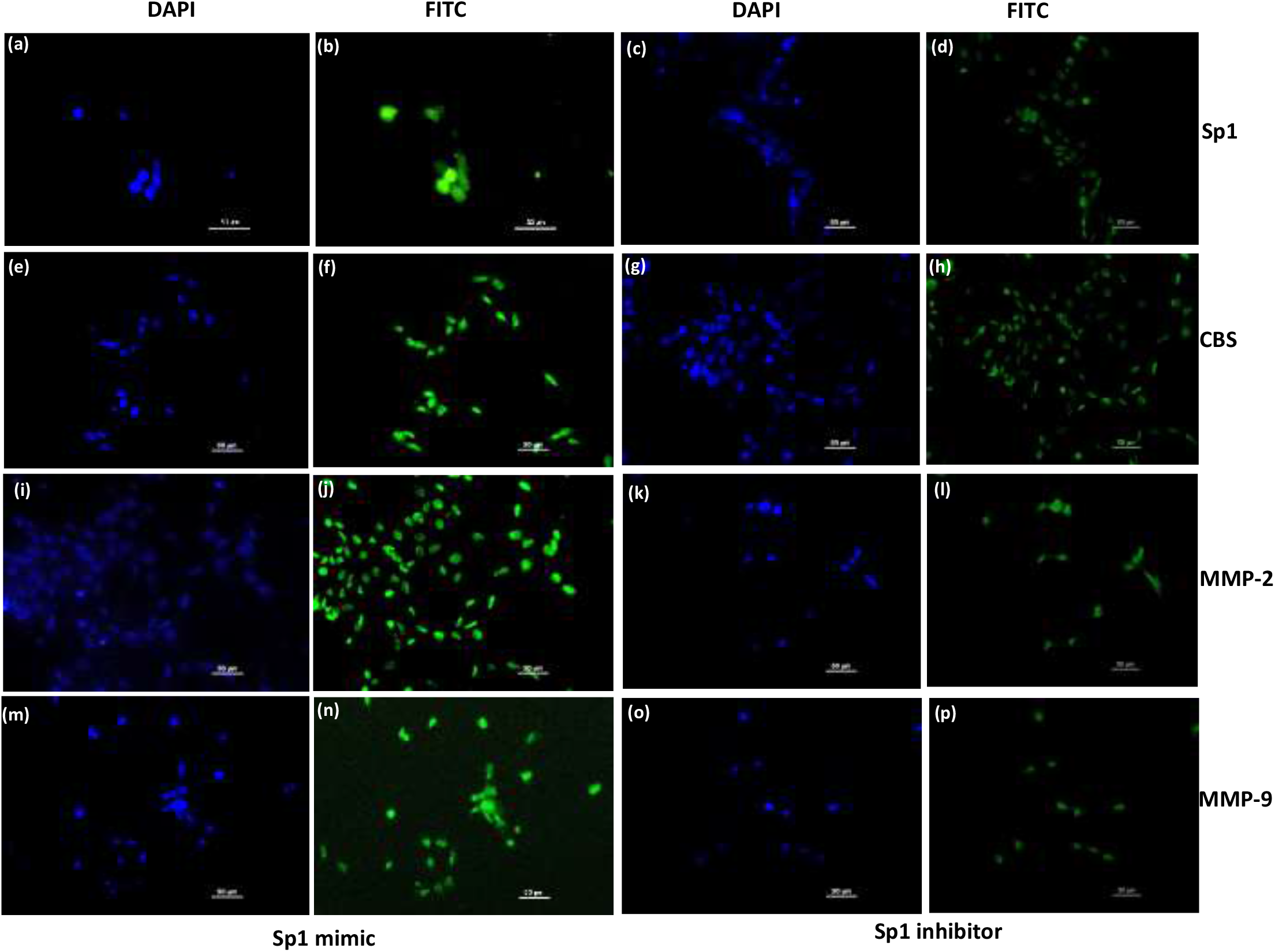
Representative Immunofluorescence images showing localization of Sp1 (b,d), CBS (f,h), MMP-2 (j,l) and MMP-9 (n,p) proteins stained with FITC when HTR-8/SVneo cells were transfected with Sp1 mimic and treated with its inhibitor. Nuclei were stained by DAPI (a,c,e,g,i,k,m,o); 20x magnification; scale bar: 50 µm.

**Figure 10:**
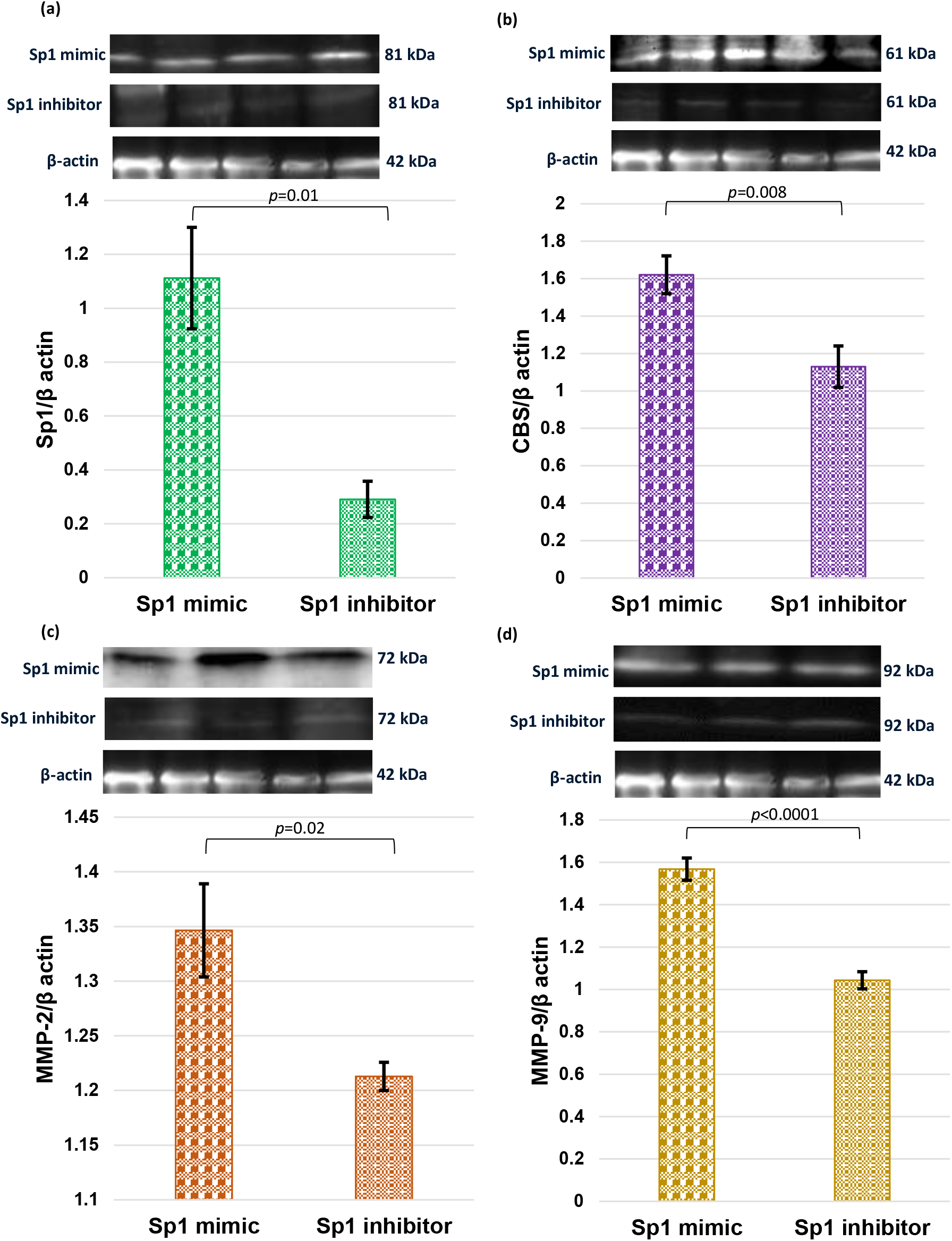
Representative images of immunoblot showing the protein expression of Sp1 (a), CBS (b), MMP-2 (c) and MMP-9 (d) when HTR-8/SVneo cells were transfected with Sp1 mimic and treated with its inhibitor. Bar diagrams represent the normalized values of Sp1 (a), CBS (b), MMP-2 (c) and MMP-9 (d) with respect to β-actin (loading control). Experiments were performed three times in triplicate (n=9). Data presented as mean ± SEM. Wilcoxon matched-pairs signed rank (a) and paired t tests (b-d) were applied.

**Figure 11.**
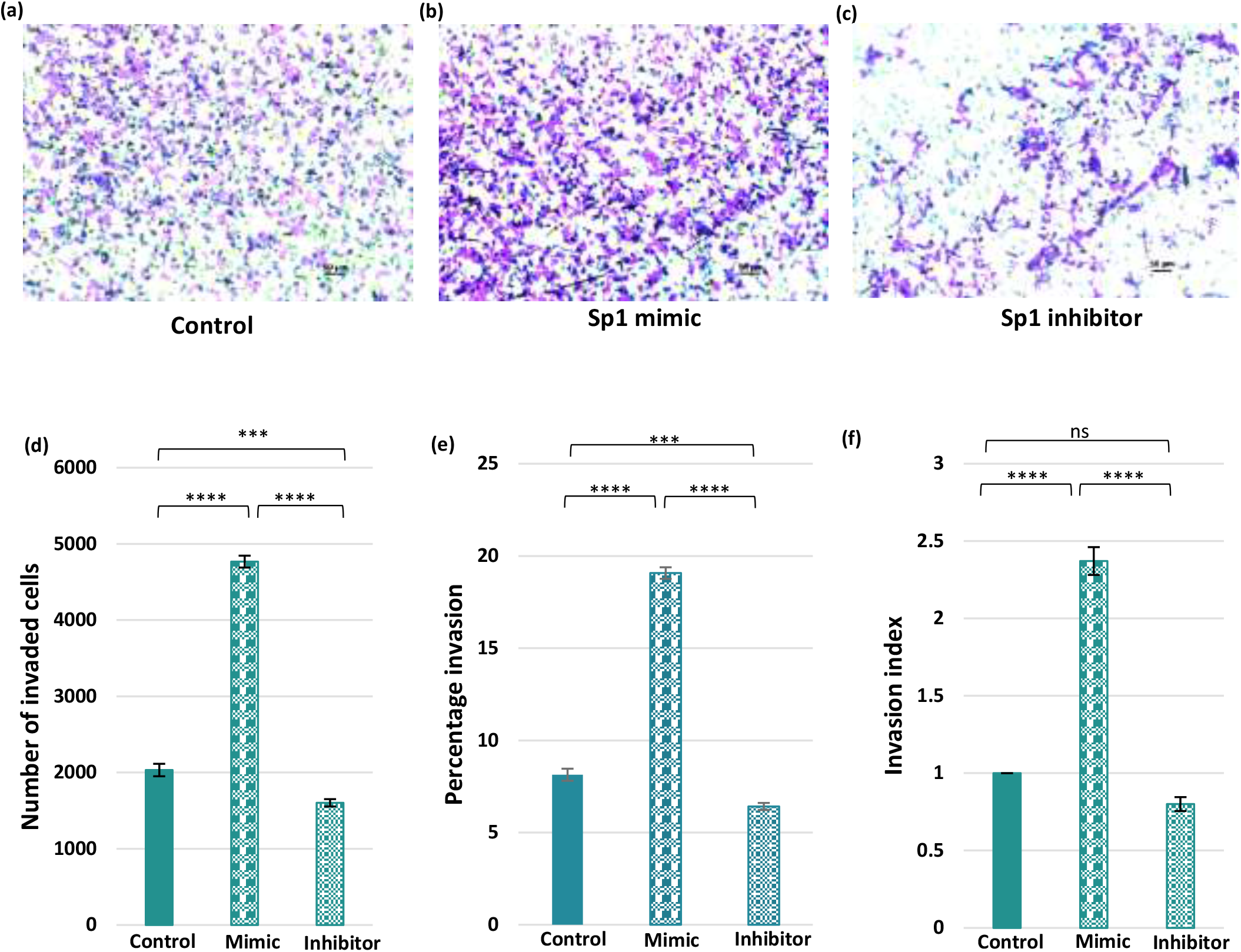
(a-c): Effect of Sp1 mimic transfection and its inhibitor treatment on the invasion capacity of HTR-8/SVneo cells. Representative matrigel transwell invasion assay images showing invaded cells on the underside of the membrane when no treatment was given (a) and after transfection with Sp1 mimic (b) and treated with its inhibitor (c), scale bar: 50 µm. 16 fields were used to count cells on each transwell insert. Bar diagrams represent number of invaded cells (d), percentage invasion (e) and invasion index (f) among various experimental groups. Experiments were performed three times in triplicate (n=9). Data presented as mean ± SEM, one way ANOVA with Bonferroni’s post hoc test was applied. ** *p*<0.01, *** *p*<0.001, \*\*\*\**p*<0.0001 ns: not significant

### Treatment of HTR-8/SVneo cells with H_2_S donor (NaHS) and CBS inhibitor (AOAA)

qRT-PCR data revealed significantly up regulated mRNA expression of CBS, MMP-2 and MMP-9 when HTR-8/SVneo cells were treated with sodium hydrogen sulphide (NaHS) whereas significant decrease in the gene expression of CBS, MMP-2 and MMP-9 was observed when the cells were treated with aminooxyacetic acid (AOAA) (Figure 12). IF staining demonstrated strong signals of CBS, MMP-2 and MMP-9 proteins when NaHS treatment was given to cells but when the cells were treated with AOAA, weak signals for CBS, MMP-2 and MMP-9 proteins were observed (Figure 13). Protein levels of CBS, MMP-2 and MMP-9 were found to be significantly up regulated when the cells were treated with NaHS however, down regulated protein levels of CBS, MMP-2 and MMP-9 were observed when the cells were subjected to AOAA treatment (Figure 14). Significant increase in the number of invaded cells, percentage invasion and invasion index were observed when the cells were treated with NaHS whereas number of invaded cells, percentage invasion and invasion index were found to be significantly decreased when AOAA treatment was given to cells (Figure 15).

**Figure 12:**
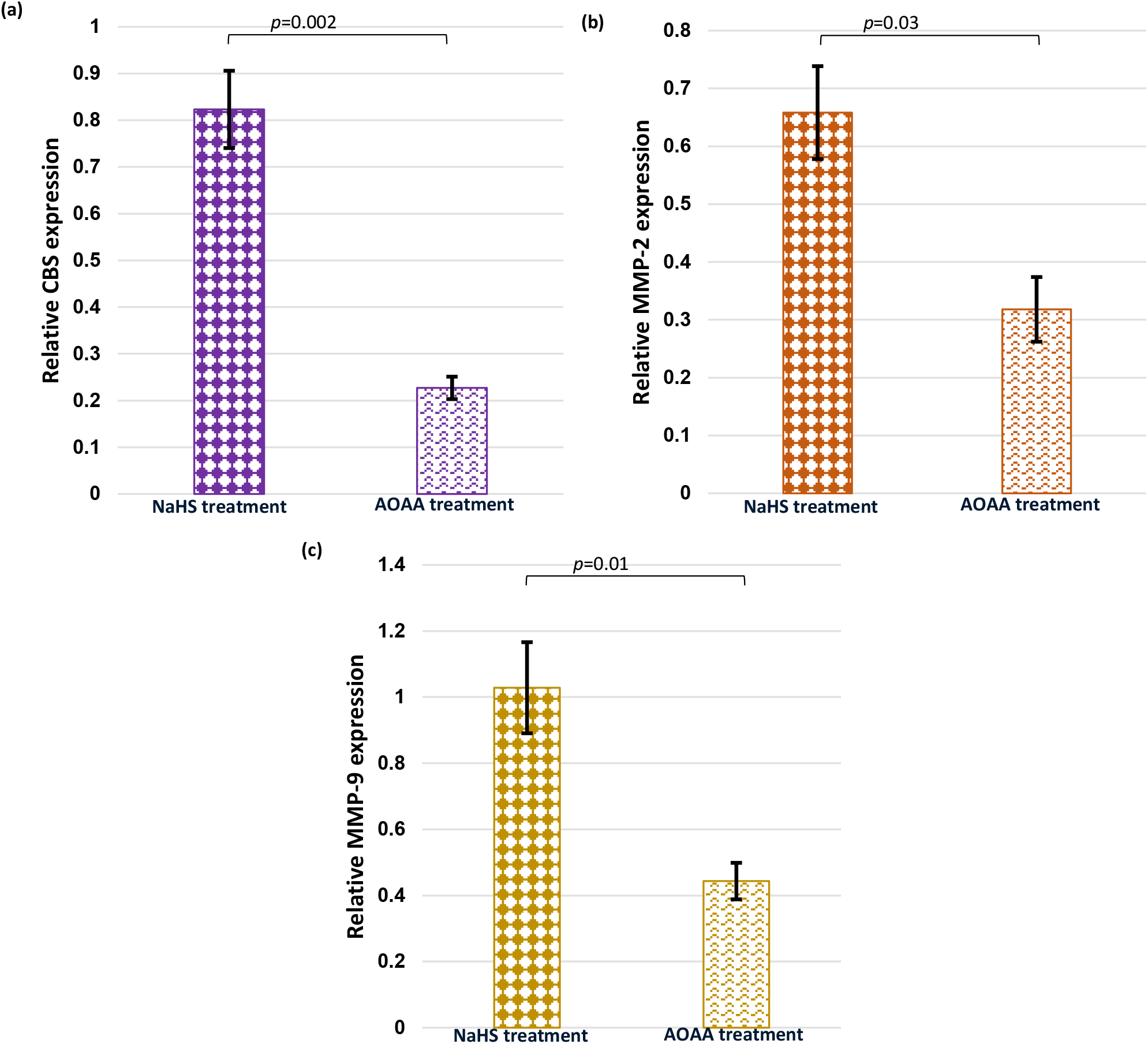
Bar diagrams represent gene expression of CBS and MMPs 2, 9 in HTR-8/SVneo cells post NaHS and AOAA treatments. Experiments were performed three times in triplicate (n=9). Data presented as mean ± SEM. Paired t (a) and wilcoxon matched-pairs signed rank tests (b,c) were applied, *p* values indicated on graphs.

**Figure 13:**
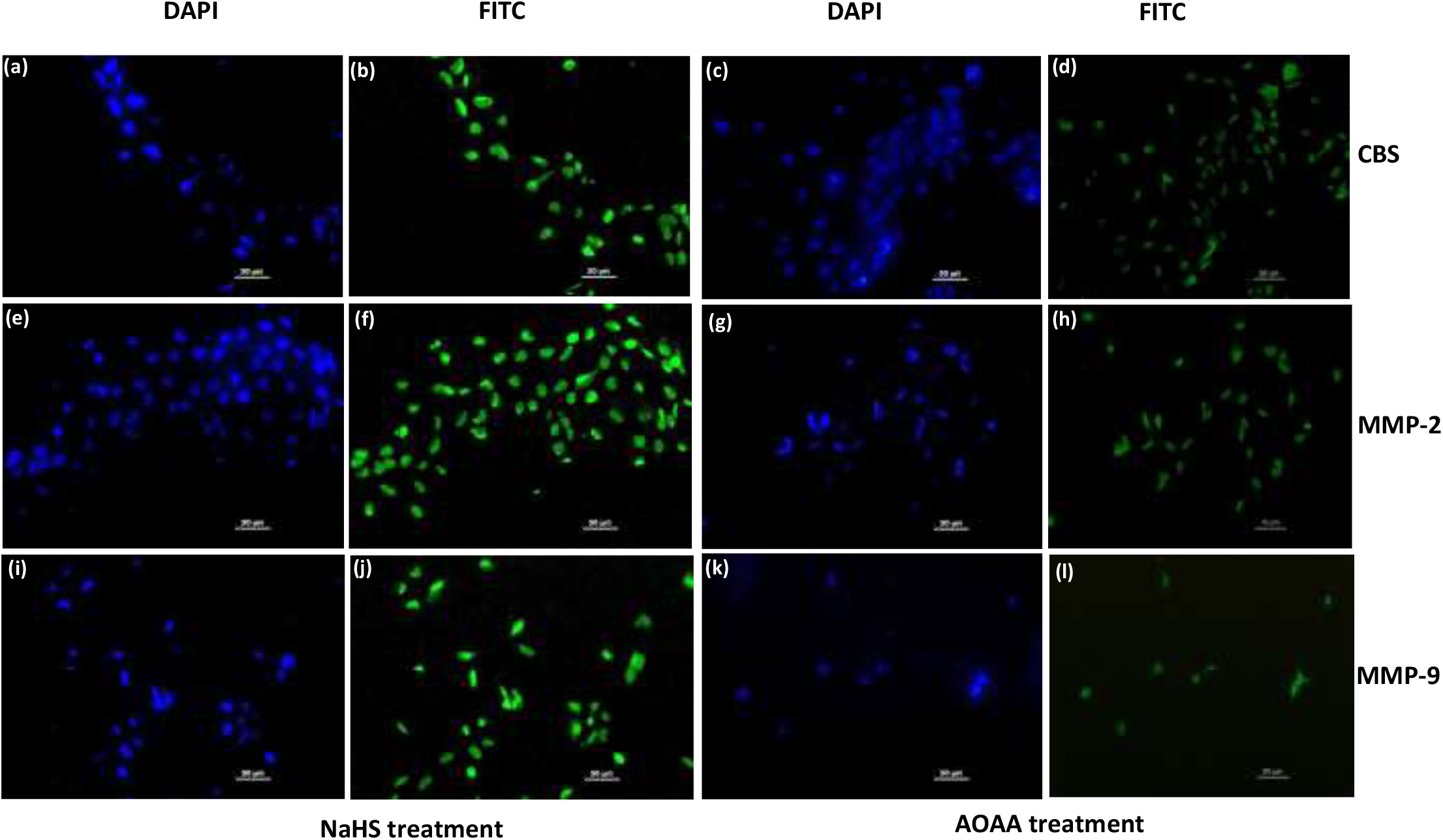
Representative Immunofluorescence images showing localization of CBS (b,d), MMP-2 (f,h) and MMP-9 (j,l) proteins stained with FITC when HTR-8/SVneo cells were treated with NaHS and AOAA. Nuclei were stained by DAPI. At 20x magnification; scale bar, 50 µm.

**Figure 14:**
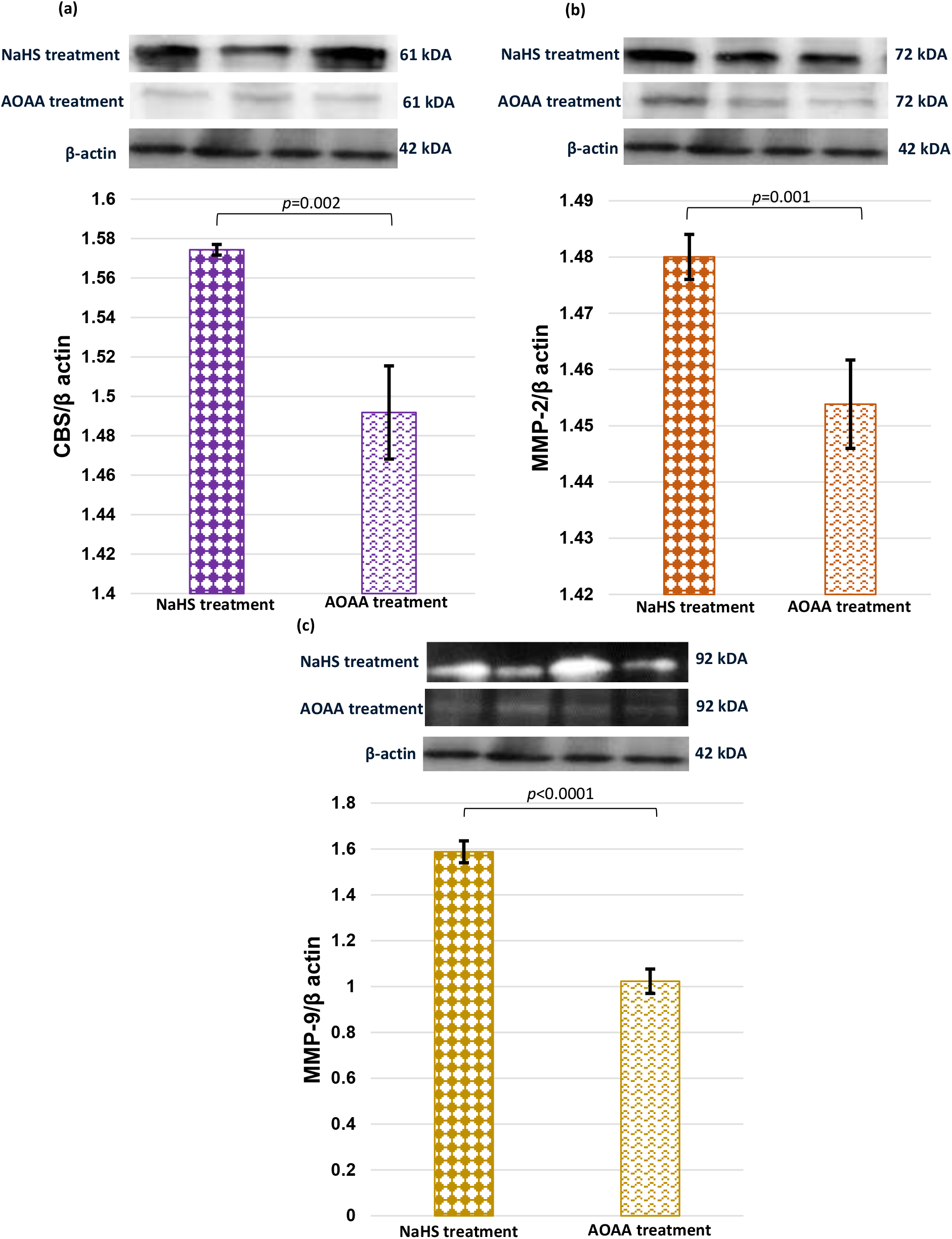
Representative images of immunoblot showing the protein expression of CBS (a), MMP-2 (b) and MMP-9 (c) when HTR-8/SVneo cells were treated with NaHS and AOAA. Bar diagrams represent the normalized values of CBS (a), MMP-2 (b) and MMP-9 (c) with respect to β-actin (loading control). Experiments were performed three times in triplicate (n=9). Data presented as mean ± SEM. Statistical analysis was done using paired t test, *p* values indicated on graphs.

**Figure 15.**
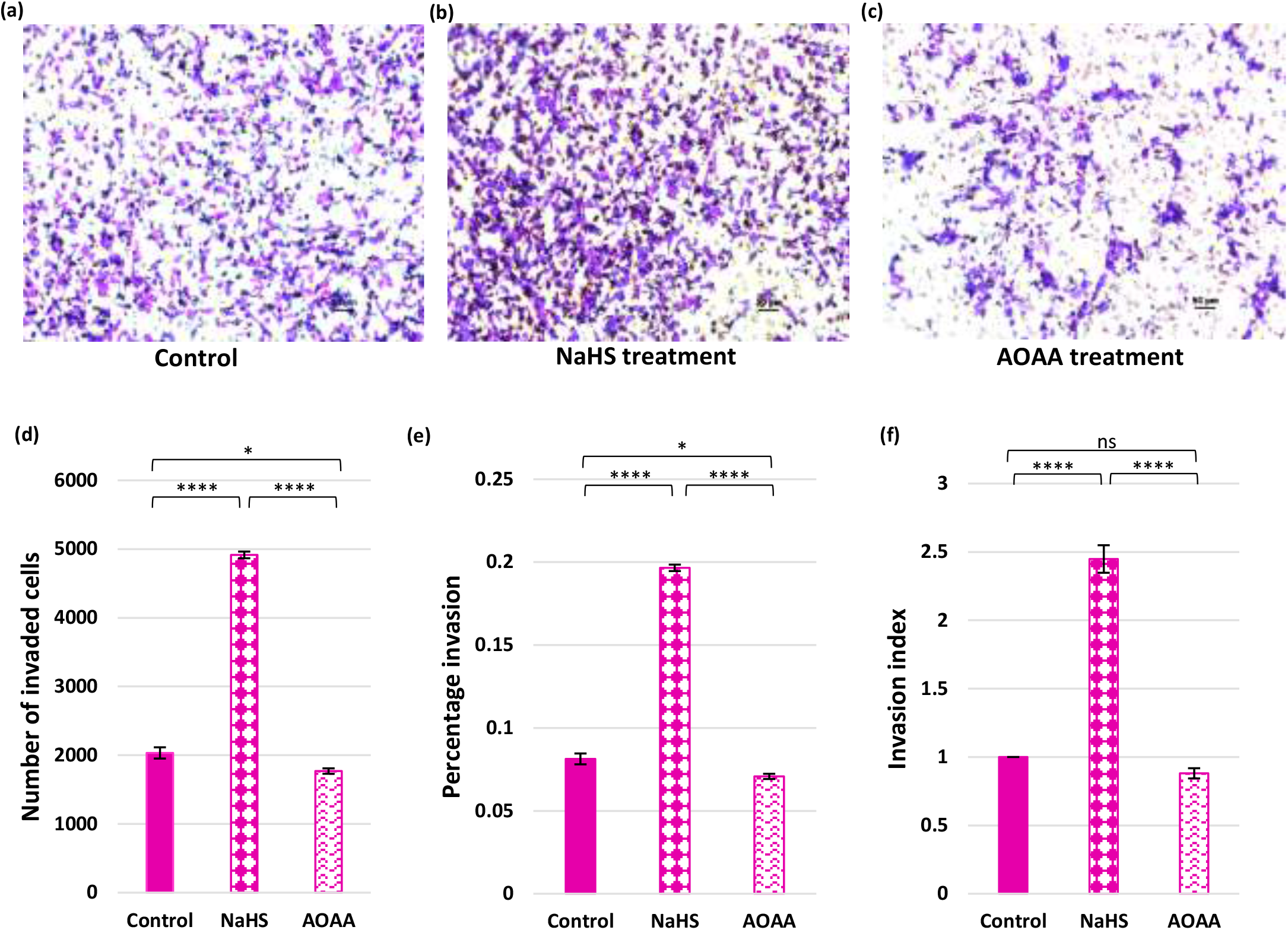
(a-c): Representative matrigel transwell invasion assay images showing invaded cells on the underside of the membrane when no treatment was given (a) and after treatment with NaHS (b) and AOAA (c), at 20x magnification; scale bar, 50 µm. 16 fields were used to count cells on each transwell insert. Bar diagrams represent number of invaded cells (e), percentage invasion (f) and invasion index (g) among various experimental groups. Experiments were performed three times in triplicate (n=9). Data presented as mean ± SEM, one way ANOVA with Bonferroni’s post hoc test was applied, *p* values indicated on graphs. \**p*<0.05, ** *p*<0.01, *** *p*<0.001, \*\*\*\**p*<0.0001 ns: not significant

## Discussion

The placenta is a transient organ during pregnancy which serves as the interface between the fetal and maternal environments and is involved in the exchange of gases, nutrients and waste products between the mother and the growing fetus^26^. Proper placental development is vital for embryo survival and health^27^ and defects in its function and development have been associated with many pregnancy complications, such as preeclampsia^28,29^, intrauterine growth restriction^30,31^, small for gestational age^32,33^, preterm birth (PTB)^3,32^ and gestational diabetes mellitus (GDM)^34^. Previous studies suggested that miRNAs are important regulators of placental development^3,4-8,32^ and have been identified in human placental tissues^9^. Also, core proteins required for miRNA biogenesis have also been detected in villous trophoblast cells^35^. In vitro studies have observed that miRNAs regulate trophoblast cell proliferation, migration, invasion, apoptosis, and angiogenesis^4,7,8,10,11^. Abnormal expression of miRNAs in placenta from women with compromised pregnancies has been reported^3,4,12,13^. Ishibashi *et al*., in 2012 have observed up regulated expression of miR-22 in the placentae from pregnancies complicated by preeclampsia as compared to control placentae by high-throughput sequencing and qRT-PCR-based array^36^. Shao *et al*. in 2017 have reported significantly enhanced miR-22 expression in early onset preeclamptic placentae as compared to their gestational age matched preterm labor controls by qRT-PCR^14^. Previous studies from our lab also observed significantly up regulated expression of miR-22 (pre miR-22 and miR-22-3p) in the early onset preeclamptic patients as compared to their gestational and maternal age matched normotensive, non-proteinuric controls. Apoptosis of *extravillous trophoblasts* was increased in early-onset preeclampsia and intrauterine growth restriction, which could in turn impair trophoblast invasion^37,38^. The invasion of spiral arteries by trophoblasts requires degrading extracellular matrix mediated by MMPs^39^ and among MMPs, MMP-2 and MMP-9 appear to play a crucial role in regulating trophoblast invasion^24^. The expression of MMP-2 and MMP-9 was down regulated in preeclamptic and IUGR placentae^25,40^. Previous studies from our lab also reported significantly down regulated expression of MMP-2 and MMP-9 in early onset preeclamptic patients as compared to healthy controls at both transcription and translation levels. Worldwide studies reported that MMP-2 and MMP-9 are targeted by many miRNAs, which may contribute to the pathogenesis of preeclampsia and intrauterine growth restriction^41-45^. Li *et al*., in 2013 showed that gene expression of MMP-2 and MMP-9 was significantly reduced upon transfection with miR-22 expression in colon cancer cells^46^ but the impact of miR-22 on the mRNA and protein expression of MMP-2 and MMP-9 in HTR-8/SVneo cells has not been reported yet. In the present study, we have transfected the cells with miR-22-3p mimic and miR-22-3p inhibitor to determine the mRNA and protein expression of MMP-2 and MMP-9 and observed significantly lower expression of MMP-2 and MMP-9 when the cells were transfected with miR-22-3p mimic, however, miR-22-3p inhibitor transfected cells had significantly higher expression of MMP-2 and MMP-9 at both transcription and translation levels. Transwell invasion assay (TIA) analysis showed significant decrease in the number of invaded cells when the cells were transfected with miR-22-3p mimic whereas miR-22-3p inhibitor transfected cells had significant increase in the number of invaded cells. We speculated that the invasion capacity of HTR-8/SVneo cells was found to be significantly decreased upon miR-22-3p mimic transfection which could be because of the significant down regulation in the expression of MMP-2 and MMP-9. Then, we studied the underlying mechanism of the regulation of MMP-2 and MMP-9 by miR-22.

Guo *et al*. in 2013 identified Specificity protein 1 (Sp1), a direct target of miR-22 and an inverse linear correlation between expression of miR-22 and Sp1 mRNA in gastric tumors was reported^15^. In the present study, we transfected the cells with miR-22-3p mimic and miR-22-3p inhibitor to determine the expression of Sp1 and observed significant down regulation in the expression of Sp1 when the cells were transfected with miR-22-3p mimic however, miR-22-3p inhibitor transfected cells had up regulated mRNA and protein expression of Sp1. Hence, we found an inverse pattern of expression between miR-22 and Sp1 in HTR-8/SVneo cells. Previous studies from our lab on clinical samples (plasma and placentae) also observed an inverse pattern of expression between miR-22 and Sp1. Huang *et al*., in 2017 reported that Sp1 bound directly to the MMP-2 promoter and increased its transcription, promoting bladder cancer cell invasion^47^. Murthy *et al*., in 2010 observed that Sp1 is one of many transcription factors that bind to the MMP-9 promoter to induce its transcription^48^. In the present study, we transfected the cells with Sp1 mimic/Sp1 expression construct and treated with Sp1 inhibitor/2,4,5 trifluoroaniline to determine the mRNA and protein expression of MMP-2 and MMP-9 and observed significantly up regulated expression of MMP-2 and MMP-9 when the cells were transfected with Sp1 mimic but 2,4,5 trifluoroaniline treatment resulted in significant down regulation in the mRNA and protein expression of MMP-2 and MMP-9. TIA analysis showed significant increase in the number of invaded cells when the cells were transfected with Sp1 expression construct whereas 2,4,5 trifluoroaniline treated cells had significant decrease in the number of invaded cells. Sp1 mimic transfection must have resulted in the significant up regulation in the levels of MMP-2 and MMP-9 which must have caused the significant increase in the invasion capacity of cells. In the light of the above findings, we speculated that miR-22 and Specificity protein-1 could regulate the mRNA and protein expression of MMP-2 and MMP-9.

Then, we studied the molecular mechanism for regulation of MMP-2 and MMP-9 by Specificity protein-1 (Sp1). Maclean *et al*. in 2004 have proposed that Sp1 has an indispensable role in the regulation of cystathionine β-synthase (CBS)^17^. CBS regulates homocysteine metabolism and contributes to hydrogen sulfide (H_2_S) biosynthesis through which it plays multifunctional roles in the regulation of cellular energetics, redox status, DNA methylation, and protein modification^49^. In their study, Maclean *et al*. have transfected Sp1 deficient fibroblast cells with an Sp1 expression construct and observed high levels of CBS expression^17^. We, in the present study have transfected the cells with Sp1 expression construct and treated with Sp1 inhibitor/2,4,5 trifluoroaniline to determine the mRNA and protein expression of CBS and observed significantly up regulated mRNA and protein expression of CBS when the cells were transfected with Sp1 mimic but when the cells were treated with Sp1 inhibitor/2,4,5 trifluoroaniline, the gene and protein expression of CBS was found to be significantly down regulated.

In the present study, for the first time, we also have proved an association between CBS and miR-22 in HTR-8/SVneo cells. Upon miR-22-3p mimic transfection in cells, we observed significant decrease in the mRNA and protein expression of CBS whereas significant increase in the mRNA and protein expression of CBS was observed when the cells were transfected with miR-22-3p inhibitor.

Subsequently, we studied the impact of CBS on the mRNA and protein expression of MMP-2 and MMP-9. Liu *et al*. in 2017 reported that exogenous NaHS (H_2_S donor) treatment significantly increased the expression levels of MMP-2 and MMP-9 in EJ cells. They treated human bladder cancer EJ cells with different concentrations of exogenous NaHS and they showed that exogenous NaHS could promote both the cell proliferation and invasion abilities^22^ by up regulating the expression of MMP-2 and MMP-9 in human bladder cancer EJ cells^22^. In the present study, we treated HTR-8/SVneo cells with NaHS and CBS inhibitor/AOAA to determine the mRNA and protein expression of MMP-2 and MMP-9 and observed significantly up regulated expression of MMP-2 and MMP-9 when the cells were treated with NaHS whereas AOAA treated cells had significantly down regulated expression of MMP-2 and MMP-9 at both transcription and translation levels. TIA analysis showed significant increase in the number of invaded cells when the cells were treated with NaHS however, significant decrease in the number of invaded cells was observed when AOAA treatment was given. When the cells were subjected to NaHS treatment, MMP-2 and MMP-9 levels must have got significantly up regulated which ultimately has led to the significant increase in the invasion capacity of cells.

## Summary

The results from the present study may point towards the role/effect of miR-22 in invasion of trophoblast cells by regulating the levels of MMP-2 and MMP-9 via Sp1 and CBS pathway. The present study for the first time will provide novel insights into the effects of miR-22, Sp1 and CBS on invasion capacity of trophoblast cells. Role of mir-22/Sp1/CBS axis in the regulation of trophoblast invasion by influencing the levels of MMP-2 and MMP-9 may become basic knowledge which ultimately contributes to investigate, clarify and prevent pregnancy related complications like preeclampsia, recurrent pregnancy loss, FGR and provide insights for pharmacological interventions in these diseases.

