## Supplementary file for "Role of MicroRNA-22 (miR-22)/Specificity protein 1 (Sp1)/Cystathionine β-synthase (CBS) axis in regulating trophoblast invasion: an in vitro study"

### Tables

| | Nucleic acid | Lipofectamine (RNA iMAX transfection reagent) | Opti-MEM |
|---|---|---|---|
| **hsa-miR-22-3p mirVana miRNA mimic** | 12.5 µl (198.5 ng/µl) | 3 µl | 150 µl |
| **MISSION Synthetic miRNA Inhibitor, Sigma** | 10 µl (238.9 ng/µl) | 3 µl | 150 µl |
| **Sp1 plasmid (Thermo)** | 2.54 µl (954.9 ng/µl) | 3 µl | 150 µl |

**Table 1: Cell transfection: Standardized doses for one reaction**

| | Concentration | Duration |
|---|---|---|
| **Sp1 inhibitor (2,4,5 Trifluoroaniline)** | 250 µM | 5 minutes |
| **CBS mimic/$H_2S$ donor (NaHS)** | 100 µM | 5 minutes |
| **CBS inhibitor (AOAA)** | 50 µM | 5 minutes |

**Table 2: Cell treatments: Standardized doses**

| Targets | Primers |
|---|---|
| **miR-22-3p** | 5’-AAGCUGCCAGUUGAAGAACUGU-3’ |
| **Precursor miR-22** | FP-GGCTGAGCCGCAGTAGTTC<br>RP-GCAGAGGGCAACAGTTCTTCAA |
| **Sp1** | FP- GGAGAGCAAAACCAGCAGAC<br>RP- AAGGTGATTGTTTGGGCTTG |
| **CBS** | FP- CTGAAGAACGAAATCCCCAA<br>RP- GCCTCCTCATCGTTGCTCTT |
| **MMP2** | FP- CGTCTGTCCCAGGATGACATC<br>RP- ATGTCAGGAGAGGCCCCATA |
| **MMP9** | FP- CGCCAGTCCACCCTTGT<br>RP- CAGCTGCCTGTCGGTGAGA |
| **U6** | FP-CTCGCTTCGGCAGCACA<br>RP-AACGCTTCACGAATTTGCGT |
| **GAPDH** | FP- AGCCGAGCCACATC<br>RP- TGAGGCTGTTGTCATACTTCTC |

**Table 3: Primers are designed by NCBI (National Centre for Biotechnology Information) and confirmed by *In silico* PCR.**
